# Population Dynamics and Resource Availability Drive Seasonal Shifts in the Consumptive and Competitive Impacts of Introduced House Mice (*Mus musculus*) on an Island Ecosystem

**DOI:** 10.1101/2022.02.23.481645

**Authors:** Michael J. Polito, Bret Robinson, Pete Warzybok, Russell W. Bradley

## Abstract

**Background:** House mice (*Mus musculus*) are widespread and invasive on many islands where they can have both direct and indirect ecological impacts on native ecological communities. Given their opportunistic, omnivorous nature the consumptive and competitive impacts of house mice on islands have the potential to vary over time in concert with resource availability and mouse population dynamics.

**Methods:** We examined the ecological niche of invasive house mice on Southeast Farallon Island, California, USA using a combination of mouse trapping, food resource surveys, and stable isotope analysis to better understand their trophic interactions with native flora and fauna.

**Results:** We found that plants were the important resource for house mice during the spring months when vegetation is abundant and mouse populations are low. However, plants decline in dietary importance throughout the summer and fall as mouse populations increase, and seabird and insect resources become relatively more available and consumed by house mice. Mouse abundance peaks and other resource availability are low in the fall months when the isotopic niches of house mice and salamanders overlap significantly indicating the potential for competition, most likely for insect prey.

**Discussion:** These results indicate how seasonal shifts in both the mouse abundance and resource availability are key factors that mediate the consumptive and competitive impacts of introduced house mice on this island ecosystem.

## Introduction

House mice (*Mus musculus*) and other rodents are some of the most widespread invasive mammals on earth; amongst vertebrates, the breadth of their global distribution is second only to that of humans (Bronson 1979; Brooke et al. 2007). In island ecosystems, house mice have been shown to have direct and indirect ecological impacts on plant, invertebrate, small mammal, and avian communities (Angel et al. 2009; Harris 2009; St Clair 2011). In addition, mice impacts can inflict substantial damage on remote islands lacking other invasive rodents such as rats (*Rattus spp*.) and/or where eradication efforts have freed mice from the constraints of competition and predation (Broome et al. 2019; Simberloff 2009). Despite this, there has been relatively less research and conservation action devoted to invasive house mice on islands, relative to other introduced mammals (Angel et al. 2009; Howald et al. 2007; Wanless et al. 2012; Wanless et al. 2007).

Southeast Farallon Island (SEFI; 37.6989° N, 123.0034° W) is located 48 km west of San Francisco off the coast of central California. This 28 ha island is a part of the Farallon Islands National Wildlife Refuge which host the largest seabird breeding colony in the contiguous United States (Johns & Warzybok 2019). SEFI also hosts an introduced house mouse population (USFWS 2013; USFWS 2019). Though the exact timing of the introduction of house mice to the South Farallon Islands is unknown, it likely occurred during the 1800’s or early 1900’s (Ainley & Lewis 1974). While early 20^th^ century data on SEFI mouse abundance are lacking, mice have had a significant presence on the islands from the late 20^th^ century to the present (Ainley & Boekelheide 1990). Closed capture modeling from a mark recapture study on SEFI provided a density estimate of 1,297 ± 224 mice per ha (95% CI: 799-1,792), one of the highest reported mouse densities for any island in the world (Grout & Griffiths 2013). Commonly, island house mouse densities range from 10 to 50 per ha (MacKay et al. 2011).

Seabirds are often particularly sensitive to invasive mammals on islands (Jones et al. 2008)). While house mice on islands are known to depredate seabird eggs and chicks and in some cases adults (Bolton et al. 2014; Dilley et al. 2015; Jones et al. 2019), there is little evidence of direct predation by mice on breeding seabirds on the South Farallon Islands. Despite over 40 years of continuous, intensive study of breeding seabirds, few depredated eggs or chicks have been detected (Ainley & Boekelheide 1990). While predation on eggs by mice can be difficult to detect in a crevice-nesting species, these observations suggest the frequency of direct seabird egg or chick predation in this population may be low (Ainley & Boekelheide 1990). Even so, there is a growing body of evidence that house mice facilitate migratory burrowing owls (*Athene cunicularia*) to overwinter on SEFI (Chandler et al. 2016; Mills 2016) which can lead to significant hyper-predation by owls on adult ashy storm petrel (*Hydrobates homochroa*) when the mouse population crashes (Nur et al. 2019).

While less studied relative to seabirds, house mice likely also have direct and indirect impacts on the ecological community on SEFI. Jones & Golightly (2006) examined the stomach contents of 57 house mice on SEFI in 2002 and 2003. They found native plants such as the maritime goldfield (*Lasthenia maritima*) constituted 63% of recovered plant material in mouse stomachs (Coulter & Irwin 2005; Jones & Golightly 2006). This contrasts with the significantly greater percentage of non-native (63-80%) plant species on SEFI, which is high given the island’s small size and relative isolation (Coulter & Irwin 2005; Hawk 2015; Jones & Golightly 2006). In addition, mouse stomachs also contained native invertebrates, including the endemic Farallon camel cricket (Jones & Golightly 2006). Even so, the interpretation of prey importance and seasonal variation in house mice diets was hampered by the inability to identify and quantify digested prey remains and an inability to sample mice between April to August (Jones & Golightly 2006). Moreover, no studies have examined the potential impacts of house mice on the endemic Farallon arboreal salamander (*Aneides lugubris farallonensis*). As salamanders feed primarily on insects and other small invertebrates (Bury & Martin 1973), it is possible that mice and salamanders compete for prey resources. Given the opportunistic, omnivorous diet of house mice (Berry 1968; Jones & Golightly 2006), it is also possible that mice consume salamander eggs when they are laid in the summer and/or small juveniles in the fall when they emerge (Boekelheide 1975).

Stable isotope analysis represents an alternative approach to traditional dietary analysis which can be used to quantify ecological niche of native and invasive species and answer ecological questions that were previously intractable (McCue et al. 2020). This approach is based on the principle that animals “are what they eat” with the carbon (δ^13^C) and nitrogen (δ^15^N) stable isotopes values of consumer tissues reflecting the abundance of these same biomarkers in their food sources (DeNiro & Epstein 1978; DeNiro & Epstein 1981). Stable isotope analysis therefore provides insights into species’ “isotopic niche”, which is analogous to the “Hutchinsonian niche” (Hutchinson 1978), as consumer tissue stable isotope values are directly influenced by what they consume (bionomic) as well as the habitat (scenopoetic) in which they live (Newsome et al. 2007). Furthermore, consumer and prey tissue stable isotope values can be incorporated into dietary mixing models to provide quantitative predictions of consumer diet compositions (Phillips et al. 2014). As consumer tissues integrate dietary information at the time of tissue synthesis it is possible to quantify consumer diets over differing time periods by examining tissues that differ in their rate of metabolic turnover (Hobson & Clark 1992; Vander Zanden et al. 2015).

The goal of our study was to use stable isotope analysis to quantitatively assess the diets of house mice on SEFI to better understand their trophic interactions with native flora and fauna ahead of their proposed eradication by the United Stated Fish and Wildlife Service (USFWS 2013). Specifically, using isotopic niche and dietary mixing model approaches we quantified seasonal shifts in the diet of house mice to determine how consumptive impacts to native bird, insect, plant, and intertidal communities vary in concert with seasonal variation in mouse population dynamics and resource availability. Furthermore, we used the isotopic niche approach to quantify the potential for consumptive and competitive interactions between introduced house mice and endemic arboreal salamanders on SEFI.

## Methods

### Study site

SEFI is the largest of the Farallon Islands, which is one of several islands that compose the Farallon Islands National Wildlife Refuge. Topographically, it is characterized by a hill that rises from the center up to 90 m high, and a wide, flat marine terrace that extends outward from the base of the hill. The terrace is widest from the southeastern to the western portions of the island, and seabirds have excavated numerous burrows within its friable soil. The hill is composed of crumbling granite cliffs that contain fissures, crevices, caves, and rocky scree fields at its base. SEFI has a temperate, maritime climate, with relatively wet winters and dry summers (average annual rainfall = 51 cm) (Kim et al. 2009). The temperature is warmest in Oct, with an average air temperature of 16.1° C, and coldest in Jan, with an average air temperature of 11.4° C (HADS; Kim et al. 2009).

Over 25% of California’s breeding marine birds, with more than 400,000 individuals of 12 species are found in the Farallon Islands National Wildlife Refuge (Johns & Warzybok 2019; Karl et al. 2001). Most areas on SEFI are occupied continually by breeding seabirds between late March and mid-August with cormorants (*Urile penicillatus, Urile pelagicus, Nannopterum auritum*) and common murres (*Uria aalge*) inhabiting rocky slopes and cliffs, storm-petrels (*Hydrobates homochroa, Hydrobates leucorhous*), auklets (*Ptychoramphus aleuticus, Cerorhinca monocerata*), pigeon guillemots (*Cepphus columba*), and tufted puffins (*Fratercula cirrhata*) nesting in rock crevices and burrows, black oystercatchers (*Haematopus bachmani*) nesting along the rocky shoreline, and Western gulls (*Larus occidentalis*) nesting across the island and most common on the flatter or more gently sloped areas (Johns & Warzybok 2019; Karl et al. 2001). Five species of pinniped visit and/or breed on SEFI including the northern elephant seal (*Mirounga angustirostris*), harbor seal (*Phoca vitulina*), Steller’s sea lion (*Eumetopias jubatus*), California sea lion (*Zalophus californianus*), and the northern fur seal (*Callorhinus ursinu*s; (Karl et al. 2001). While hoary bats (*Aeorestes cinereus*) and Mexican free-tail bats (*Tadarida brasiliensis*) have been recorded visiting the island, house mice are the only breeding terrestrial mammals present on SEFI (Karl et al. 2001; Tenaza 1966). The islands flora includes at least 44 species, 26 of which are non-native (Coulter & Irwin 2005). Honda et al. (2017) reported 11 insect orders representing 60 families, 107 genera and 112 insect species identified on SEFI. Two endemic taxa are found on SEFI: the Farallon camel cricket (Rentz 1972) and the Farallon arboreal salamander (Van Denburgh 1905).

### Ethics statement

Sampling was approved by the United States Fish and Wildlife Service under a cooperative agreement with Point Blue Conservation Science and the California Department of Fish and Game scientific (permit no. SC–008556). All vertebrate sampling protocols were approved by and adhered to statutes of the Institutional Animal Care and Use Committee of the Woods Hole Oceanographic Institution (permit no. 4855-001).

### Mouse abundance and resource availability

We created an index of mice abundance based on monthly trapping success on 4 transect lines spread across available habitats at SEFI (Irwin 2004; Nur et al. 2019). Trapping was conducted for each of 3 nights per month between March 2001 and March 2004, and again from December 2010 to March 2012, and finally September 2016 through January 2018. All sampling periods used the same transects, each with 7 traps per transect. For the 2010-2012 and 2016-2018 effort, 5 additional traps were added; these incorporated more of the vertical aspect of the island topography. Trapping efforts used D-Con^®^ Ultra Set^®^ covered snap traps baited with peanut butter and oats. Trapping success was determined as the proportion of trap-nights set per monthly session (either 84 [2001-2004] or 99 [2010-2012 and 2016-2018]) in which house mice were captured.

Precipitation data (inches of rain) was collected on SEFI using a standard US National Weather Service rain gauge (∼1 m elevation) at the same location at noon Pacific Standard Time every day throughout the time series examined here. Precipitation data was used as a proxy for both terrestrial environmental conditions and vegetation phenology as the majority of vegetation on SEFI senesces or dies during the summer and recovers in the winter and spring when seasonal rainfall begins (Coulter & Irwin 2005).

Insect density (indiv./m^2^) on SEFI was evaluated during nine collecting trips between February 2013 and October 2014. Each trip lasted roughly 12 continuous days. Methods of insect collection and density estimates are detailed in Honda et al. (2017).

We used daily year-round assessments of adult seabird carcasses found during routine island operations as proxy of seabird resource availability to house mice on SEFI. As operations and research protocols have remained standard throughout the time series, effort is relatively consistent throughout the period. Adult birds had their primary tips clipped with a knife to prevent double counting. For the purpose of this analysis, we used carcass count numbers from one common burrow nesting species whose breeding habitat overlapped mouse sampling areas, Cassin’s Auklet (*Ptychoramphus aleuticus*), as an indicator of overall seabird resource abundance.

To obtain an estimate of the seasonal variability in the abundance (indiv./month) of Farallon Arboreal Salamanders we obtained data from a long-term monitoring program of this species on SEFI where standardized cover boards were used to assess salamander populations (Lee et al. 2012).

The mouse trapping success and resource availability datasets described above were compiled between 2001 and 2017 unless noted otherwise below. For all metrics, monthly values were compiled and averaged to obtain seasonal values (i.e. Spring, Summer, Fall, Winter) for each year that data were available (Table S1). Average values were calculated for salamander abundance, mouse trapping success, and insect density. Total monthly values were compiled for rainfall and bird carcasses. We used Kruskal Wallis Test with non-parametric post-hoc comparison to test seasonal differences in mouse trapping success and the availability of food resources.

### Tissue sampling

We sampled muscle and liver tissues from house mice (n= 63 individual) during three different seasons in the spring (April), summer (August), and fall (October) in 2013 that reflect the natural oscillation of mouse abundance on SEFI (Irwin 2006). Mouse liver tissue stable isotope values reflects short term diet on the scale of days to weeks, while muscle tissues stable isotope values reflect diets on the scale of weeks to months (DeMots et al. 2010). During the spring (April) and fall (October) sampling period in 2013 we also collected representative samples of prey items such as muscle tissue from intertidal snails, vegetative tissue from common plants, and whole-bodied insects (Table S2). We also collected seabird tissues (i.e. muscle, egg membrane, guano, etc.) during the summer (August) once these resources became readily available on the island (Table S2). Finally, we sampled tail clips from arboreal salamanders found under cover boards in the spring (April; n = 16) and fall (October; n = 16) when they are most active on the island. All sampled were stored frozen (−20°C) prior to processing for stable isotope analysis.

### Stable isotope analysis

All samples were freeze-dried and then homogenized using a mortar and pestle. Dried homogenized muscle, liver and tail clip samples underwent lipid extraction using 2:1 chloroform:methanol. Samples were placed in a glass vail with a solvent volume 10 timesLgreater thanLsample volume and sonicated in a water bath for 15 min and then decanted. This procedure was repeated for a total of three cycles and the samples were rinsed in DI water and oven dried at 60°C for 24 h to remove any remaining solvent. We flash-combusted (PDZ Europa ANCA-GSL elemental analyzer) approximately 1.0 mg of each animal tissue sample and 3.0 mg of each plant tissue sample loaded into tin cups to analyze for carbon and nitrogen isotopes (δ^13^C and δ^15^N) through interfaced PDZ Europa 20-20 continuous-flow stable isotope ratio mass spectrometers (CFIRMS). Raw δ values were normalized using glutamic acid (G-17; G-9/USGS-41), bovine liver (G-13), peach leaves (G-7) and nylon 5(G-18) as standard reference materials. Sample precision based on repeated standard reference materials was 0.1‰ for both δ^13^C and δ^15^N. Stable isotope ratios are expressed in δ notation in per mil units (‰), according to the following equation:

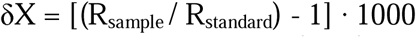

Where X is ^13^C or ^15^N and R is the corresponding ratio ^13^C /^12^C or ^15^N /^14^N. The R_standard_ values were based on the Vienna PeeDee Belemnite (VPDB) for δ^13^C and atmospheric N_2_ for δ^15^N.

### Isotopic niche analysis

We assessed variation in isotopic niche (Newsome et al. 2007) position, width, and overlap across seasons (i.e. spring, summer, and fall) and tissue (muscle vs. liver) in house mice and between species (i.e. house mice and arboreal salamander) within each season using both multivariate and univariate techniques. We compared isotopic niche positions by computing the Euclidean distance (ED) between group centroids (δ^13^C and δ^15^N bivariate means) following the methods of Turner et al. (2010). Isotopic niche positions were considered to be different if the ED between species examined was significantly greater than zero after comparison to null distributions generated by a residual permutation procedure. If niche positions differed, we then examined the results of univariate general linear models (GLM) and Tukey-Kramer multiple comparison tests to determine which axis (δ^13^C and/or δ^15^N) contributed to niche differences across seasons or between species (Hammerschlag-Peyer et al. 2011). To examine individual consistency in the isotopic niche of house mice we tested for relationships between individual’s liver (i.e. shorter-term dietary signal) and muscle (i.e. longer-term diet signal) stable isotope values using Pearson correlations.

In addition, we explore variation in niche area and overlaps using standard ellipse areas which can be interpreted as a measure of the core isotopic niche of a population (Jackson et al. 2011). We calculated the Bayesian estimation of standard ellipse area (SEA_b_) for each group to compare 2-dimensional niche areas of house mice and arboreal salamanders among species and seasons. We then used the resulting Bayesian posterior probability distributions of SEA_b_ estimates in pairwise-tests to identify significant differences in SEA_b_ at the P < 0.5 level. Lastly, we compared isotopic niche overlap between groups calculated as the proportion of standard ellipse area corrected for sample size (Jackson et al. 2011) for each group that overlap with a comparison group’s standard ellipse area (SEA_c_ overlap). Among other things, this isotopic niche overlap approach provided us with a quantitative estimate of the potential for competitive overlap between mice and salamanders.

### Dietary mixing model analyses

We used univariate (δ^13^C or δ^15^N), GLM and Tukey-Kramer multiple comparison tests to examine differences across major prey resources (seabirds, intertidal snails, plants, and insects) collected in each season. As not all prey resources (i.e. seabirds) were collected in every season we grouped prey resources by taxa and season into a single factor when conducting GLM analyses. We then used a Bayesian mixing model (Parnell & Inger 2016; Parnell et al. 2010) in the R environment (Ver. 3.6.2) to quantify the relative use of these four major prey resources by house mice in each season. This model estimates the probability distributions of multiple source contributions to a mixture while accounting for the observed variability in source and mixture isotopic signatures, elemental concentration, and dietary isotopic fractionation. We focused our mixing model analyses on liver tissues given the strong correlation in isotopic values found between mouse tissues (see Results below) and the more constrained and shorter isotopic turnover time in liver (DeMots et al. 2010; MacAvoy et al. 2005). Separate models were run in each season (spring, summer, summer). As the discriminatory power of Bayesian mixing models can decline markedly above six or seven prey sources (Phillips et al. 2014), we *a priori* defined five statistically, ecologically, and taxonomically relevant prey resource groups for incorporation into our mixing model analysis. As little to no differences in prey resource stable isotopes values were observed across seasons (see Results below), we used the same prey resource δ^13^C and δ^15^N values averaged across all seasons in each model analyses. As there is no existing evidence to support or refute the possibility that house mice may also consume arboreal salamanders on SEFI we also explored a separate subset of mixing models that include arboreal salamanders as a prey resource for house mice. We used diet to consumer isotopic discrimination factors for liver tissue (δ^15^N: +4.3±0.2; δ^13^C: +0.7±0.3) derived from captive studies of house mice on a controlled diet of wheat and corn (Arneson & MacAvoy 2005). We incorporated elemental concentration dependence (Phillips & Koch 2002) into the model, and ran 1 million iterations, thinned by 15, with an initial discard of the first 40,000 resulting in 64,000 posterior draws.

## Results

### Seasonal trends in mouse abundance and resource availability

Mouse trapping success varied by season (H_3_=19.1058, p<0.001) with lower trapping success in the spring relative to the late summer and fall (Fig. 1). These long-term trends broadly concurred with the trapping success observed in 2013, when mouse tissue and food resource samples were collected for stable isotope analyses (Table S1, Fig. 1). Precipitation also differed across seasons (H_3_=43.8296, p<0.001) being lowest in the summer, intermediate in the spring and fall, and highest in the winter (Fig. 1). Insect density varied significantly across seasons (H_3_=20.2933, p<0.001) with lower values in the summer and higher values in the spring and winter (Fig. 1). Seabird carcass abundance also varied significantly across seasons (H_3_=47.3676, p<0.001) with higher values in spring and summer relative to fall and winter (Fig. 1). Finally, salamander abundances differed across seasons (H_3_=30.3015, p<0.001) with higher average counts in spring and winter relative to summer (Fig. 1).

**Figure 1.**
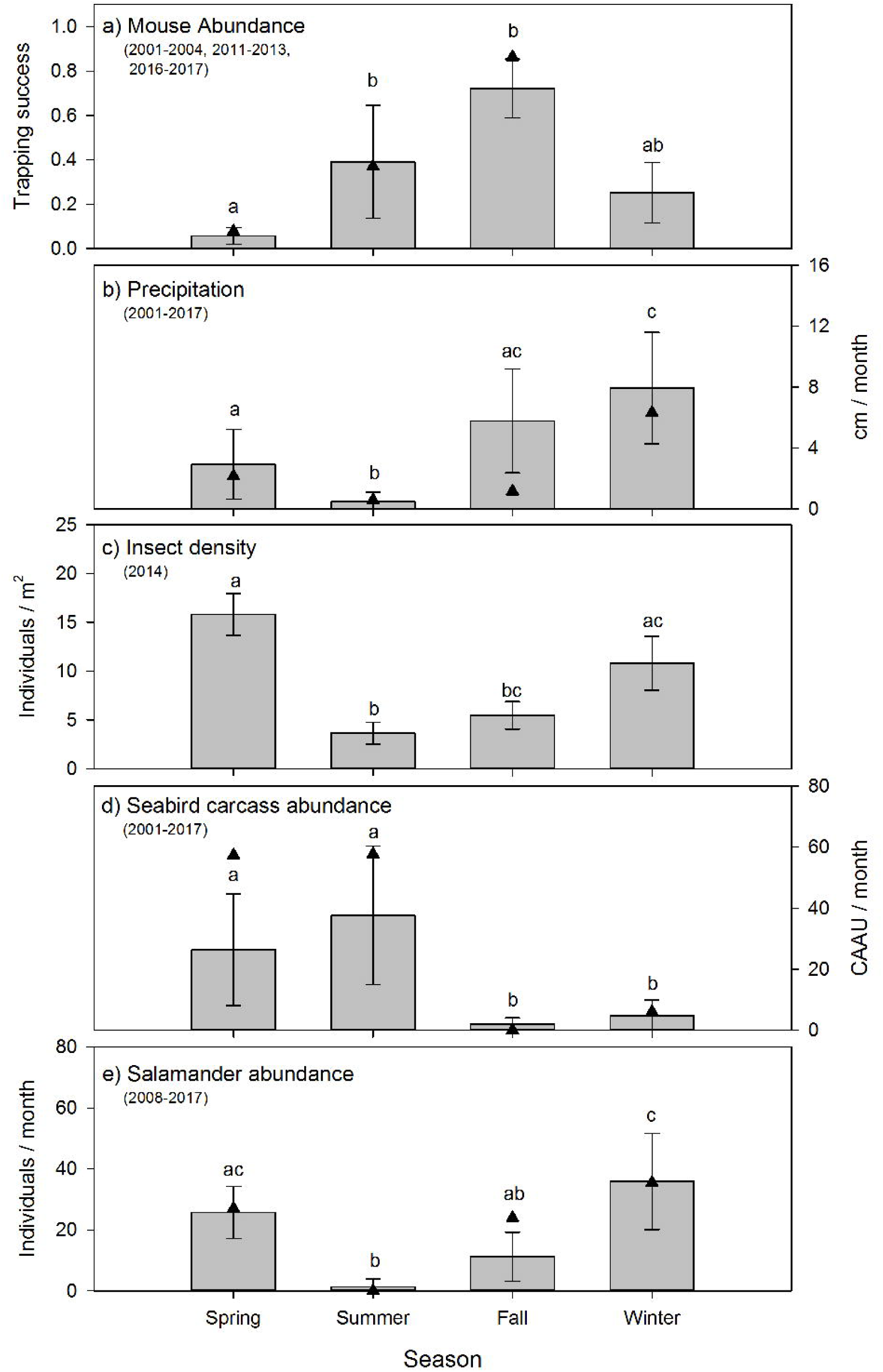
The relative abundance of a) invasive house mice (*Mus musculus*), b) monthly precipitation amount, c) seasonal insect densities, d) monthly seabird carcass abundance (CAAU: Cassin’s auklet (*Ptychoramphus aleuticus*), and e) monthly arboreal salamanders (*Aneides lugubris farallonensis*) abundance on Southeast Farallon Island, CA during the spring, summer, fall and winter seasons for the years listed. Triangles represent the seasonal mean values in 2013 when tissues were collected for stable isotope analyses. Grey bar and whiskers represent seasonal mean (±SD) values across all of the years listed for each dataset. For each dataset, seasons that share a superscript are not significantly different at the P < 0.05 level.

### Isotopic niche of house mice and arboreal salamanders

The isotopic niche position of house mice differed significantly in all pairwise comparisons across seasons for both liver (ED = 4.01-5.37‰, p < 0.001) and muscle (ED = 2.00-5.02‰, p < 0.030) tissues. House mice stable isotope values differed among seasons (δ^13^C: F_2,126_ = 66.61, p < 0.001; δ^15^N: F_2,126_ = 26.60, p < 0.001) but not by tissue type (δ^13^C: F_1,126_ = 66.61, p = 0.878; δ^15^N: F_1,126_ = 0.92, p = 0.339) or the interactions between these two factors (δ^13^C: F_1,126_ = 2.32, p = 0.681; δ^15^N: F_1,126_ = 1.22, p = 0.299). Seasonal differences in the isotopic niche position of house mice were due to lower tissue δ^13^C values in the spring relative to the summer and fall as well as lower tissue δ^15^N values in the summer relative to the spring (liver and muscle) and fall (liver only; Table 1). Muscle and liver tissue stable isotope values were positively correlated with one another for both δ^13^C (r = 0.908, p <0.001) and δ^15^N (r = 0.946, p <0.001) values.

**Table 1.** The carbon (δ^13^C) and nitrogen (δ^15^N) stable isotope values and C/N ratio of house mice (*Mus musculus*) and endemic arboreal salamanders (*Aneides lugubris farallonensis*) during the the spring, summer and fall sampling periods on Southeast Farallon Island, CA in 2013. Groups that share a superscript are not significantly different at the P < 0.05 level.

| Taxa, Tissue | Season | *n* | C/N | $\delta^{13}\text{C}$ (‰) | $\delta^{15}\text{N}$ (‰) |
| --- | --- | --- | --- | --- | --- |
| House mice, liver | Spring | 23 | 3.7±0.1 | -24.5±0.6<sup>a</sup> | 27.0±1.8<sup>a</sup> |
|  | Summer | 20 | 3.5±0.1 | -22.0±1.3<sup>b</sup> | 22.2±4.2<sup>b</sup> |
|  | Fall | 20 | 3.5±0.1 | -21.2±1.6<sup>bc</sup> | 26.1±2.3<sup>a</sup> |
| House mice, muscle | Spring | 23 | 3.4±0.1 | -24.4±1.1<sup>a</sup> | 26.8±1.5<sup>a</sup> |
|  | Summer | 20 | 3.4±0.2 | -21.8±1.6<sup>b</sup> | 22.5±4.6<sup>b</sup> |
|  | Fall | 20 | 3.4±0.1 | -21.7±1.4<sup>b</sup> | 24.5±1.8<sup>ab</sup> |
| Arboreal salamander, tail clip | Spring | 16 | 3.4±0.2 | -21.5±1.0<sup>b</sup> | 25.9±1.4<sup>a</sup> |
|  | Fall | 16 | 3.3±0.1 | -20.0±1.4<sup>c</sup> | 24.5±1.5<sup>ab</sup> |

The isotopic niche area of house mice differed significantly across seasons due to smaller SEA_b_ values in spring relative to summer and fall for liver tissues and higher SEA_b_ values in summer relative to spring and fall for muscle tissues (Table 2). While the isotopic niche of house mice did not overlap among seasons when examined using liver tissues, niche overlap based on muscle tissue was higher between spring and summer (7.6-18.9%) and spring and fall (11.0-12.4%) relative to fall and summer (0.0-0.1%; Fig 1.).

**Table 2.** Isotopic niche area of house mice (*Mus musculus*) and endemic arboreal salamanders (*Aneides lugubris farallonensis*) during the spring, summer and fall sampling periods on Southeast Farallon Island, CA in 2013. Isotopic niche area of mice and salamanders are presented as Bayesian estimation of standard ellipse area (SEA_b_) and their corresponding 95% credibility intervals. Groups that share a superscript are not significantly different at the P < 0.05 level.

| Taxa, tissue | Season | *n* | Mean SEA<sub>b</sub> (‰<sup>2</sup>) | SEA<sub>b</sub> 95% CI (‰<sup>2</sup>) |
| --- | --- | --- | --- | --- |
| House mice, liver | Spring | 23 | 4.05<sup>a</sup> | 2.86-5.61 |
|  | Summer | 20 | 9.40<sup>bc</sup> | 6.46-13.39 |
|  | Fall | 20 | 9.54<sup>bc</sup> | 6.58-13.51 |
| House mice, muscle | Spring | 23 | 5.20<sup>ab</sup> | 3.67-7.25 |
|  | Summer | 20 | 12.87<sup>c</sup> | 8.90-18.22 |
|  | Fall | 20 | 5.88<sup>ab</sup> | 4.05-8.42 |
| Arboreal salamander, tail clip | Spring | 16 | 4.37<sup>a</sup> | 2.89-6.44 |
|  | Fall | 16 | 6.52<sup>abc</sup> | 4.29-9.66 |

The isotopic niche position of house mice and arboreal salamanders differed significantly in all within-season, pairwise comparisons of arboreal salamander tail clips and mouse liver (spring: ED = 3.19‰, p < 0.001; fall: ED = 2.02‰, p = 0.004) and muscle (spring: ED = 3.05‰, p < 0.001; fall: ED = 1.71‰, p = 0.011) tissues. This was due to higher δ^13^C values in arboreal salamanders’ tail clips relative to mouse liver and muscle tissues in the spring (F_2,118_ = 63.61, p < 0.001) and muscle tissues only in the fall (F_2,118_ =6.03, p = 0.004; table 1). In contrast, arboreal salamanders tail clip δ^15^N values did not differ from house mouse liver and muscle tissues in the spring (F_2,118_ = 2.23, p = 0.101) or the fall (F_2,118_ = 3.40, p = 0.057; table 1).

The isotopic niche width of arboreal salamanders did not differ significantly between seasons, nor did they differ from estimates of house mice niche area calculated from liver or muscle tissues within each season (Table 2). Isotopic niche overlap between spring and fall for arboreal salamanders ranged from 16.0-24.9%. The isotopic niche of arboreal salamanders in the spring did not overlap with those of house mice estimated using either liver or muscle tissues (Fig. 2). In contrast, the isotopic niche of arboreal salamanders in the fall showed significant overlap with the isotopic niche area (SEAc) of house mice estimated using liver tissue (16.3%) or muscle tissues (52.3%) (Fig. 2).

**Figure 2.**
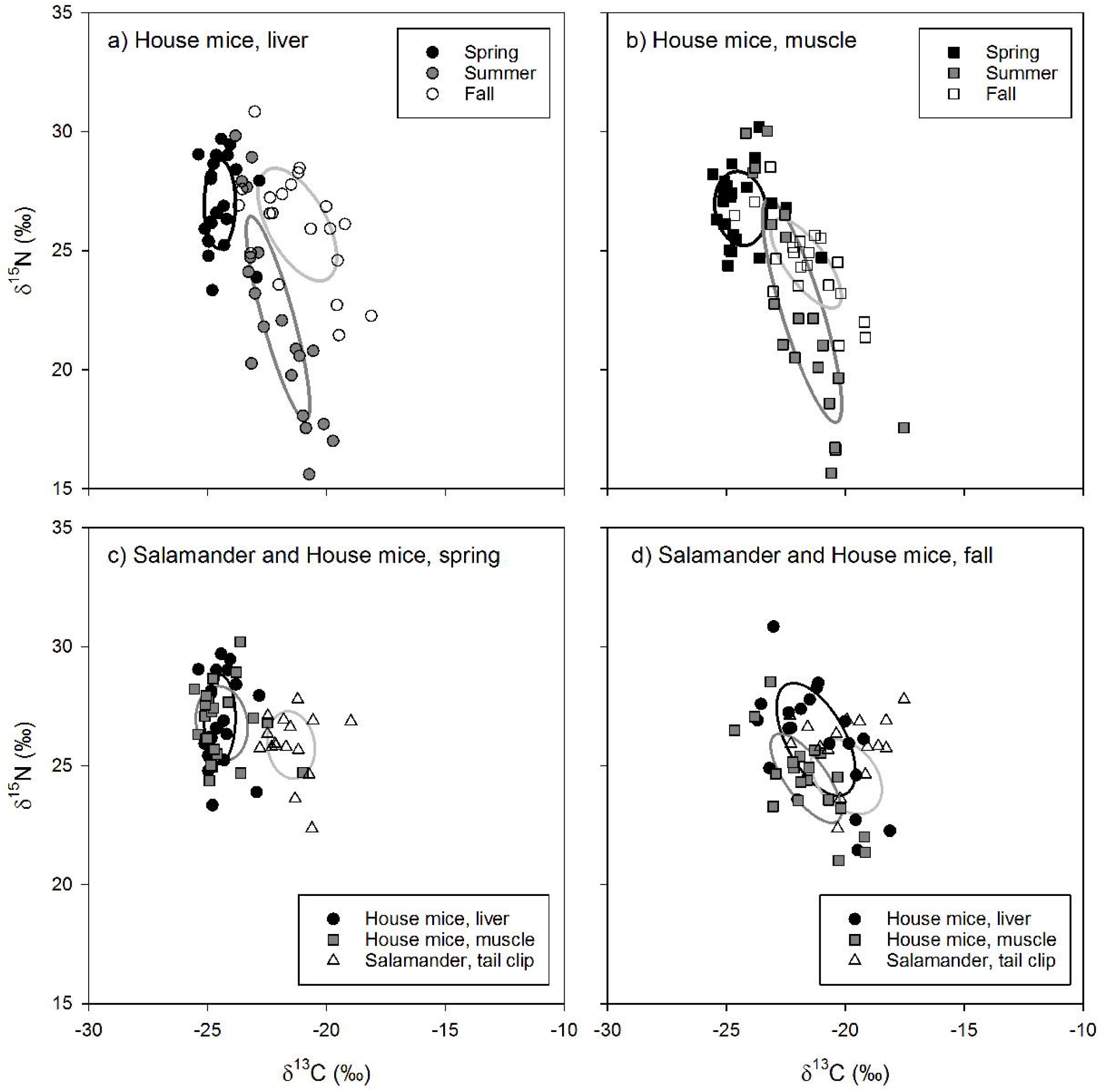
The carbon (δ^13^C) and nitrogen (δ^15^N) stable isotope values and isotopic niche areas of house mice (*Mus musculus*) liver and muscle tissues and endemic arboreal salamander (*Aneides lugubris farallonensis*) tail clip tissues during the spring, summer and fall sampling periods on Southeast Farallon Island, CA in 2013. Isotopic niche areas are presented a standard ellipse area corrected for sample size (SEA_c_; Jackson et al. 2011).

### Dietary mixing model analyses

Differences in prey resource stable isotope values between major taxonomic groupings were more common than differences within major taxonomic groupings collected in different seasons for both δ^13^C (F_4,152_ = 132.60, p < 0.001) and δ^15^N (F_4,152_ = 38.25, p < 0.001) values. For example, plant and intertidal resource δ^13^C and δ^15^N values, and insect and arboreal salamander δ^15^N values did not differ between seasons (Table 1, Table S1). Insect and arboreal salamander δ^13^C values differed slightly between seasons, though these differences were small (1.5-1.9‰) relative to the differences observed between major taxonomic groupings (2.2-16.5‰; Table S1, Table S2). Because of these findings, and because not all prey resources were collected in each season even though they were available, prey resource δ^13^C and δ^15^N values were averaged across all seasons by major taxonomic groupings for subsequent analyses (Table S2). These averaged prey resource δ^13^C (F_4,152_ = 237.27, p < 0.001) and δ^15^N (F_4,152_ = 75.35, p < 0.001) values differed significantly among major taxonomic groups thus providing isotopically unique endmembers for incorporation into the isotopic mixing model (Fig. 3).

**Figure 3.**
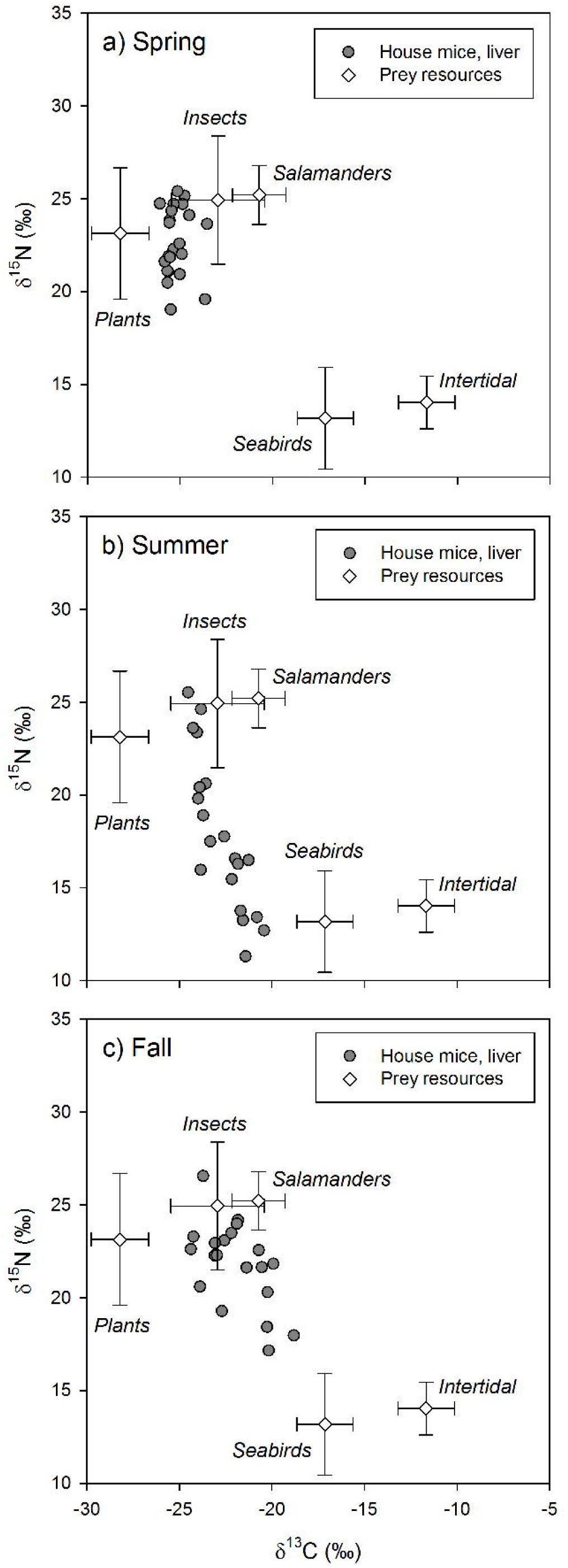
The carbon (δ^13^C) and nitrogen (δ^15^N) stable isotope values of house mice (*Mus musculus*) liver tissues and possible prey resources in the spring, summer and fall sampling periods on Southeast Farallon Island, CA in 2013. House mice stable isotope values been adjusted by subtracting dietary isotopic discrimination factors for liver (δ^15^N: 4.3±0.2; δ^13^C: 0.7±0.3) tissues (Arneson & MacAvoy 2005).

When examined without arboreal salamanders as a possible prey resource, stable isotope mixing model analysis indicate seasonal shifts in the diet composition of house mice on SEFI. While there is overlap in 95% credibility intervals between some groups, plants dominated the diet in the spring, followed by a smaller proportion of insects, and very little intertidal and seabird resources (Table 3). The importance of plants decreased in the summer, while the relative importance of seabirds during this time increased to the highest observed across all seasons (Table 3). Fall diets were characterized by a relatively increased importance of insect and to a lesser extent intertidal resources, a continued decline in the importance of plants, and a decrease in the importance of seabirds. When prey resources were aggregated *a posteriori*, there were clear differences in the relative use of marine (i.e., seabirds and intertidal) vs. terrestrial (i.e., plants and insects) resources by mice across seasons. While terrestrial resources were more important to mice in all three seasons, the relative importance of marine resources were higher in the summer and fall, relative to the spring (Table 3).

**Table 3.**
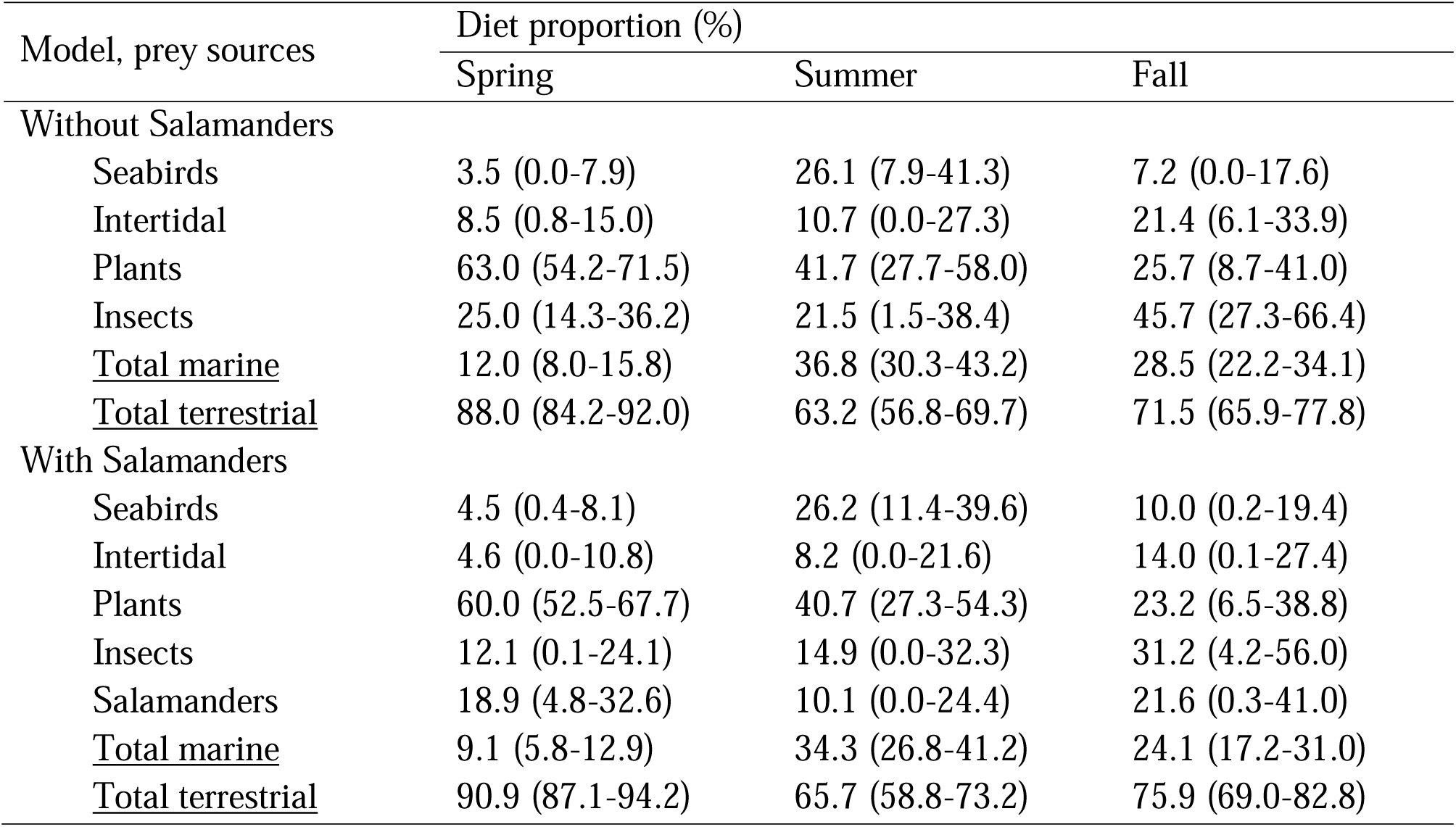
The predicted diet composition of house mice (*Mus musculus*) in the spring, summer and fall sampling periods on Southeast Farallon Island from a stable isotope-based mixing model both with and without endemic arboreal salamanders (*Aneides lugubris farallonensis*) as possible prey resources. Diet proportions for each prey resource individually, and grouped into marine (seabirds and intertidal) and terrestrial (insects, plants, and salamanders) prey resources, are presented as mean values and 95% creditability intervals in parentheses.

| Model, prey sources | Diet proportion (%) |  |  |
| --- | --- | --- | --- |
|  | Spring | Summer | Fall |
| Without Salamanders |  |  |  |
| Seabirds | 3.5 (0.0-7.9) | 26.1 (7.9-41.3) | 7.2 (0.0-17.6) |
| Intertidal | 8.5 (0.8-15.0) | 10.7 (0.0-27.3) | 21.4 (6.1-33.9) |
| Plants | 63.0 (54.2-71.5) | 41.7 (27.7-58.0) | 25.7 (8.7-41.0) |
| Insects | 25.0 (14.3-36.2) | 21.5 (1.5-38.4) | 45.7 (27.3-66.4) |
| <u>Total marine</u> | 12.0 (8.0-15.8) | 36.8 (30.3-43.2) | 28.5 (22.2-34.1) |
| <u>Total terrestrial</u> | 88.0 (84.2-92.0) | 63.2 (56.8-69.7) | 71.5 (65.9-77.8) |
| With Salamanders |  |  |  |
| Seabirds | 4.5 (0.4-8.1) | 26.2 (11.4-39.6) | 10.0 (0.2-19.4) |
| Intertidal | 4.6 (0.0-10.8) | 8.2 (0.0-21.6) | 14.0 (0.1-27.4) |
| Plants | 60.0 (52.5-67.7) | 40.7 (27.3-54.3) | 23.2 (6.5-38.8) |
| Insects | 12.1 (0.1-24.1) | 14.9 (0.0-32.3) | 31.2 (4.2-56.0) |
| Salamanders | 18.9 (4.8-32.6) | 10.1 (0.0-24.4) | 21.6 (0.3-41.0) |
| <u>Total marine</u> | 9.1 (5.8-12.9) | 34.3 (26.8-41.2) | 24.1 (17.2-31.0) |
| <u>Total terrestrial</u> | 90.9 (87.1-94.2) | 65.7 (58.8-73.2) | 75.9 (69.0-82.8) |

These same seasonal trends are also apparent when including salamanders as a possible prey resource. Plants are most important in the spring and decline in contribution to the diet throughout the summer and fall (Table 3). Seabird resources are consumed relatively more in the summer and insect resources are relatively more important in the fall. Arboreal salamanders are predicted to contribute to between 10.1-21.6% of house mice diet (Table 3). Mixing models that include Arboreal salamanders as a possible prey resource tend to predict lower insect contribution to house mice diets than those that do not include salamanders likely due to the fact that these two groups have the most similar stable isotope values of the possible prey resources examined in our study (Table 3, Table S2). When prey resources were aggregated *a posteriori* into marine vs. terrestrial resources, the results of models that included salamanders did not differ from those that did not (Table 3).

## Discussion

Our study provides insights into the diet of invasive house mice on Southeast Farallon Island (SEFI). Specifically, the use of stable carbon and nitrogen isotope analysis combined with isotopic niche and dietary mixing model approaches allowed us to quantitatively assess the diets and foraging niches of house mice on Southeast Farallon Island to better understand their interactions with native flora and fauna. We found that plants are the most important resource for house mice in the spring when plants are most abundant and house mouse populations are low. However, plants decline in importance throughout the summer and fall as mouse populations increase, and seabird and insect resources become relatively more available and important to house mice. In addition, when the mouse population is high on SEFI and possibly other resources are less abundant, the isotopic niches of house mice and salamanders overlap significantly indicating the potential for competition, most likely for insect prey. These results indicate how seasonal shifts in both the mouse population and resource availability are key factors that drive the consumptive and competitive impacts of introduced house mice on this island ecosystem.

Our results broadly agree with dietary studies of invasive mice on other island ecosystems. On Antipodes Island, the predatory and competitive impacts of house mice vary with seasons, tracking resource availability from abundant invertebrates and land birds over summer to terrestrial vegetation and seabirds in winter (Russell et al. 2020). Similarly, Bicknell et al. (2020) found that the diets of introduced St Kilda field mouse (*Apodemus sylvaticus hirtensis*) on Hirta varied among sub-populations that had differing spatial access to seabird resources, and between the seabird breeding and non-breeding seasons. Smith et al. (2002) observed both habitat and season specific variation in the diet of invasive house mice on Marion Island, with plant material most important in summer and invertebrates most important in winter and spring. Furthermore, climate change is predicted to exacerbate the direct and indirect impacts of house mice on Marion Island via shifts in mouse abundance, primary productivity, decomposition, and nutrient cycling (Smith 2002). The impacts of house mice on SEFI are also likely to be affected by climate change, which is predicted to increase in temperatures, shifts rainfall patterns, lead to changes in the phenology and composition of island vegetation, and impact invertebrate communities (USFWS 2019). These results and others highlight the highly plastic, omnivorous dietary niche of mice inhabiting island ecosystems and their ability to quickly response to seasonal and spatial resource pulses (Drever et al. 2000; Le Roux et al. 2002; Quillfeldt et al. 2008; Shiels et al. 2013).

While our study used stable isotope analysis, a prior study of house mice diet on SEFI relied solely on stomach content analyses (Jones & Golightly 2006). They found that plants and insects were found in mouse stomach throughout the year, but that eggshell and feather fragments were only recovered in the summer months (Jones & Golightly 2006). However, stomach contents reflect only a “snapshot” of an individual’s recent diet. Therefore, rodent stomach contents can often be highly variable, biased towards prey that does not readily digest, and may underestimate the amount of soft-bodied prey (Hansson 1970; Jordan 2005; Monadjem 1997). In addition, prey recovered from the stomach contents are often unable to be identified (Jordan 2005). Because the stable isotope values of mouse tissues integrate dietary information over days to months, they avoid many of the digestive and temporal biases inherent to stomach content analyses (DeMots et al. 2010; MacAvoy et al. 2005). Even so, the stable isotope-based analyses performed in our study broadly agrees with the seasonal dietary trends in frequency of occurrence observed by Jones & Golightly (2006), while in some cases also providing more constrained estimates of the relative dietary proportion of each prey source. Moreover, the positive correlation between mouse liver and muscle stable isotope values found in our study suggest a degree of individual consistency in mouse diets over time (Bearhop et al. 2004; Herman et al. 2017).

Stable isotope analysis, like stomach content analysis, cannot discern whether the seabird resources consumed by mice was the result of predation or scavenging. During the summer seabird breeding season on SEFI there is ample opportunity for invasive mice to scavenge chick and adult carcasses, lost or deserted eggs, seabird regurgitate, and/or guano. Studies on other islands using camera traps, behavioral observations, and other methods have found clear evidence of mice actively predating seabird egg and live chicks (Bicknell et al. 2020; Wanless et al. 2012; Wanless et al. 2007). The little observational data that exist on SEFI suggest the potential for a low degree of direct seabird egg or chick predation (Ainley & Boekelheide 1990). However, with the information currently available it is not possible to fully assess the relative occurrence of seabird predation vs. scavenging by house mice on SEFI.

Our stable isotope-based mixing model estimates of mouse diets on SEFI have additional assumptions and limitations. For example, we parametrized the mixing model using isotopic discrimination factors for liver tissue derived from captive studies of house mice on a controlled diet (Arneson & MacAvoy 2005). If the isotopic differences between diets and mouse livers on SEFI varied from those measured in this controlled study, they could bias the resulting estimations of proportion the proportional contribution of each diet component (Bond & Diamond 2011). However, given that these discrimination factors are tissue and species-specific, and that following their incorporation mouse tissue isotopic values were well within the range of the isotopic values of dietary resource selected based on prior knowledge there is sufficient support for their applicability in our study (Phillips et al. 2014). Moreover, while deviations from assumed discrimination factors might lead to variation in predicted absolute dietary proportions, they would be unlikely to alter our conclusions on seasonal shifts in the relative importance in dietary resource among seasons. This is because we observed clear shifts in mouse tissue δ^13^C and δ^15^N values that made them more, or less, similar to the isotopic values of specific dietary resources in each season (Fig. 3).

In addition, stable isotope analysis cannot conclusively determine if mice on SEFI are consuming endemic arboreal salamander. This approach only provides their possible dietary contribution if an *a priori* assumption is made that mice on SEFI do consume salamanders. In addition, given the similar stable isotope values of salamanders and insects the stable isotope mixing model used in these analyses had difficulties distinguishing the relative importance of these two resources to mouse diets when both resources were included in the analyses (Table 3). Therefore, while our mixing models results indicate that it is possible that mice do consume salamanders on SEFI, it is not possible to establish this definitively. Similarly, the isotopic similarity of native vs. non-native plants as well as camel crickets vs. other insects on SEFI precludes the ability of stable isotope based mixing models to identify the relative dietary contributions between these groups of mouse food resources (Table S2).

Introduced house mice may also act as competitors to native species in island ecosystems. Russell et al. (2020), found mice had a strong impact on snipe (*Coenocorypha aucklandica meinertzhagena*e) through resource competition. Specifically, during the summer snipe feed significantly lower in the food chain due to competition with mice for invertebrates, which may prevent snipe reaching adequate breeding condition (Russell et al. 2020). While mice have been shown to have an indirect impact on adult ashy storm petrel populations via hyper-predation by owls on SEFI (Nur et al. 2019), no studies to date have explored the potential for resource competition among mice and other native fauna on SEFI.

Our study used an isotopic niche approach to compare the trophic niches of mice and salamanders on SEFI to gain insights into the possibility of competitive interactions between these two species. This approach is based on using δ^13^C and δ_15_N values in bi-variate space as a proxy of the n-dimensional niche space defined by Hutchinson (1959) to represent an animal’s scenopoetic (e.g. basal carbon / habitat use) and bionomic (e.g. trophic level / diets) niche axes, respectively (Newsome et al. 2007). In this context, isotopic niche areas metrics described the range of dietary resources and habitats used by mice and salamanders, while isotopic niche overlap metrics provided a proxy measure of the potential for competitive overlap between mice and salamanders.

The isotopic niche of mice and salamanders did not overlap with salamanders in the spring but did exhibit overlap with salamanders in the fall. This isotopic overlap between species suggests the potential for resource competition in the fall when mouse populations are at their highest (Fig.1) and mixing model results suggest mice are consuming a relatively higher proportion of insect resources (Table 3). During the same time insect densities and salamander abundance are low following peak densities and counts in the winter and spring (Fig.1). Insects and other invertebrates are an important component of arboreal salamanders’ diets (Bury & Martin 1973). While it is not possible to know if lower insect densities are limiting salamander abundances at this time, given their isotopic overlap with mice the potential for resource competition exists. Even so, differences in isotopic turn-over rates, metabolic factors, and/or sub-habitat scale spatial variation in stable isotope baselines among taxa has the potential to lead to isotopic niche overlap even when species differ in diets (HetteLTronquart 2019). Therefore, future studies seeking to quantify the diets and trophic niche overlap of mice and salamanders at SEFI would benefit from integrating stable isotope analyses with other dietary proxies such as micro-analyses of gut contents and DNA metabarcoding analyses.

## Conclusions

In conclusion, our study highlights the omnivorous and opportunistic diets of house mice on SEFI. We found that mouse diets on SEFI rapidly respond to seasonal shifts in resource availability. Plants are the most important dietary resource in the spring, the importance of seabirds is highest in the summer, and insects become relatively more important in the fall. In addition, during the fall months when mouse numbers are highest the isotopic niches of house mice and salamanders overlap significantly indicating the potential for competition, most likely for insect prey.

The trophic flexibility of house mice, in combination with dramatic seasonal shifts in this invasive consumer’s overall abundance, drive the consumptive and competitive impacts of introduced house mice on this island ecosystem.

## Acknowledgements

We thank Farallon Islands National Wildlife Refuge managers and staff, and Farallon Island biologists who collected data over the past decades. Research on South Farallon Islands was made possible by a cooperative agreement with the United States Fish and Wildlife Service. Additional funding was provided by the Bently Foundation, Elinor Patterson Baker Trust, Marisla Foundation, Giles W. and Elise G. Mead Foundation, Frank A. Campini Foundation, Bernice Barbour Foundation, Kimball Foundation, RHE Charitable Foundation, Volgenau Foundation, and individual donors. The Farallon Patrol provided logistical support. This is Point Blue Contribution number 2172.

## Notes

### Competing Interest Statement

The authors have declared no competing interest.

