## Supplemental Table 1 for "Population Dynamics and Resource Availability Drive Seasonal Shifts in the Consumptive and Competitive Impacts of Introduced House Mice (*Mus musculus*) on an Island Ecosystem"

**Table S1**. The mean±SD house mice (*Mus musculus*) trapping success (%), precipitation (cm/month), insect density (indiv./$m^2$), seabird (Cassin's auklet *Ptychoramphus aleuticus*) carcass abundance (indiv./month), arboreal salamanders (*Aneides lugubris farallonensis*) abundance (indiv./month) on Southeast Farallon Island, CA during the winter (Dec., Jan., Feb.), spring (Mar., Apr., May), summer (Jun., Jul., Aug.), and fall (Sep., Oct., Nov.) seasons from 2001 to 2018. SD are not provided when less than three months of data were available in a specific year and season.

| Year | Season | Mouse trapping | Precipitation | Insect | Seabird | Salamander |
|---|---|---|---|---|---|---|
| 2001 | Winter | | 2.9±3.3 | | | |
| 2001 | Spring | 0.09±0.06 | 0.3±0.5 | | 12.0±11.5 | |
| 2001 | Summer | 0.29±0.07 | 0.1±0.2 | | 40.7±47.7 | |
| 2001 | Fall | 0.78±0.06 | 4.8±3.7 | | | |
| 2002 | Winter | 0.31±0.33 | 1.9±0.4 | | | |
| 2002 | Spring | 0.02±0.03 | 0.7±1.0 | | 67.3±62 | |
| 2002 | Summer | 0.12±0.13 | 0.1±0.1 | | 68.0±43.4 | |
| 2002 | Fall | 0.55±0.26 | 4.3±4.6 | | 4.3±2.1 | |
| 2003 | Winter | 0.37±0.27 | 1.8±0.5 | | 5.0±1.0 | |
| 2003 | Spring | 0.03±0.04 | 1.5±1.8 | | 56.7±34.1 | |
| 2003 | Summer | 0.24±0.21 | 0.1±0.1 | | 95.3±47.6 | |
| 2003 | Fall | 0.60±0.16 | 3.2±3.6 | | 0.3±0.6 | |
| 2004 | Winter | 0.22±0.18 | 5.0±3.7 | | 2.3±4.0 | |
| 2004 | Spring | 0.11 | 1.3±1.8 | | 29.0±19.1 | |
| 2004 | Summer | | 0.0±0.0 | | 53.0±36.9 | |
| 2004 | Fall | | 1.7±2.2 | | 0.0±0.0 | |
| 2005 | Winter | | 4.8±1.2 | | 1.0±1.7 | |
| 2005 | Spring | | 3.1±1.7 | | 9.0±11.5 | |
| 2005 | Summer | | 0.2±0.2 | | 6.0±5.6 | |
| 2005 | Fall | | 4.4±3.3 | | 0.3±0.6 | |
| 2006 | Winter | | 5.1±2.5 | | 1.3±0.6 | |
| 2006 | Spring | | 0.1±0.2 | | 13.0±5.6 | |
| 2006 | Summer | | 0.0±0.0 | | 8.0±3.5 | |
| 2006 | Fall | | 2.0±1.0 | | 0.0±0.0 | |
| 2007 | Winter | | 1.8±1.1 | | 0.3±0.6 | |
| 2007 | Spring | | 0.1±0.1 | | 9.0±3.6 | |
| 2007 | Summer | | 0.6±0.7 | | 32.7±29.1 | |
| 2007 | Fall | | 3.3±2.6 | | 0.0±0.0 | |
| 2008 | Winter | | 1.6±2.1 | | 0.7±0.6 | 74.8±16.6 |
| 2008 | Spring | | 0.1±0.1 | | 20.7±11.0 | 21.7±35.8 |
| 2008 | Summer | | 0.1±0.1 | | 43.3±14.6 | 0.0±0.0 |

**Table S1 (continued)**

| Year | Season | Mouse trapping | Precipitation | Insect | Seabird | Salamander |
|---|---|---|---|---|---|---|
| 2008 | Fall | | 1.3±0.5 | | 1.3±2.3 | 19.5 |
| 2009 | Winter | | 2.6±2.6 | | 0.7±1.2 | 35.2±1.3 |
| 2009 | Spring | | 0.4±0.6 | | 15.0±10.5 | 22.2±33.4 |
| 2009 | Summer | | 0.7±0.8 | | 15.0±7.9 | 0.0±0.0 |
| 2009 | Fall | | 2.5±2.6 | | 2.7±2.5 | 17.5 |
| 2010 | Winter | | 2.9±1.4 | | 1.7±2.9 | 42.5±27.8 |
| 2010 | Spring | | 1.1±0.9 | | 11.7±2.5 | 41.9±20.9 |
| 2010 | Summer | | 0.0±0.0 | | 14.3±15.6 | 7.8±6.7 |
| 2010 | Fall | | 1.9±1.6 | | 0.0±0.0 | 13.0±5.3 |
| 2011 | Winter | 0.46±0.33 | 3.3±1.7 | | 2.3±2.1 | 43.3±19 |
| 2011 | Spring | 0.08±0.01 | 2.3±2.8 | | 18.0±17.3 | 19.2±21.8 |
| 2011 | Summer | 0.48±0.29 | 0.8±1.1 | | 32.0±6.6 | 4.5±2.6 |
| 2011 | Fall | 0.74±0.36 | 1.4±1.2 | | 2.3±2.1 | 12.8±16.6 |
| 2012 | Winter | 0.09±0.05 | 1.4±1.1 | | 2.7±3.1 | 35.5±21.5 |
| 2012 | Spring | 0.0±0.0 | 2.0±2.2 | | 20.7±2.5 | 36.0±27.8 |
| 2012 | Summer | | 0.1±0.1 | | 45.0±11.3 | 0.7±1.2 |
| 2012 | Fall | | 2.4±2.6 | | 5.3±6.1 | 14.0±8.5 |
| 2013 | Winter | | 2.5±3.1 | | 6.0±8.7 | 35.4±12.7 |
| 2013 | Spring | 0.08 | 0.9±0.5 | | 57.3±21.4 | 27.0±23.6 |
| 2013 | Summer | 0.37 | 0.2±0.2 | | 57.7±50.3 | 0.0±0.0 |
| 2013 | Fall | 0.86 | 0.5±0.5 | | 0.0±0.0 | 24.0±8.5 |
| 2014 | Winter | | 1.9±3.3 | 3.2±0.8 | 21.3±18.7 | 26±36.2 |
| 2014 | Spring | | 1.4±1.5 | 4.7±0.6 | 41.0±26.0 | 32.5±23.0 |
| 2014 | Summer | | 0.1±0.1 | 1.1±0.3 | 42.0±31.4 | 0.0±0.0 |
| 2014 | Fall | | 1.4±1.4 | 1.6±0.4 | 5.0±2.6 | 3.0±4.2 |
| 2015 | Winter | | 4.8±6.8 | | 9.7±9.1 | 38.2±36.8 |
| 2015 | Spring | | 0.4±0.5 | | 27.7±16.9 | 16.3±24.5 |
| 2015 | Summer | | 0.1±0.1 | | 25.0±18.7 | 0.0±0.0 |
| 2015 | Fall | | 0.5±0.8 | | 1.0±1.7 | 5.5 |
| 2016 | Winter | | 3.1±2.3 | | 6.0±2.6 | 30.5±14.3 |
| 2016 | Spring | | 2.1±2.6 | | 14.3±8.4 | 22.7±21.5 |
| 2016 | Summer | | 0.1±0.0 | | 31.3±24.2 | 0.0±0.0 |
| 2016 | Fall | 0.63±0.01 | 2.4±2.3 | | 3.3±3.5 | 1.0±1.4 |
| 2017 | Winter | 0.16±0.03 | 6.2±1.9 | | 7.3±6.4 | 17.7±12.7 |
| 2017 | Spring | 0.06 | 2.0±1.8 | | 26.3±4.9 | 17.2±16.4 |
| 2017 | Summer | 0.85 | 0.1±0.1 | | 30.7±14 | 0.0±0.0 |
| 2017 | Fall | 0.89±0.01 | 0.7±1.0 | | 5.3±5.1 | 2.0±3.5 |
| 2018 | Winter | 0.15±0.06 | 2.7±3.6 | | 6.5±2.1 | 15.8±9.5 |
