## Supplemental Table 2 for "Population Dynamics and Resource Availability Drive Seasonal Shifts in the Consumptive and Competitive Impacts of Introduced House Mice (*Mus musculus*) on an Island Ecosystem"

**Table S2**. The carbon ($\delta^{13}C$) and nitrogen ($\delta^{15}N$) stable isotope values and C/N ratio of potential prey resources collected on Southeast Farallon Island, CA in 2013. Major taxonomic groups that share a superscript are not significantly different at the P < 0.05 level.

| Group | Taxa | Tissue | Season | *n* | C:N | $\delta^{13}C$ (‰) | $\delta^{15}N$ (‰) |
|---|---|---|---|---|---|---|---|
| Plant | *Lasthenia maritima* | Vegetation | Spring | 6 | 10.9±0.8 | -28.5±1.4 | 26.0±2.4 |
| | | | Fall | 6 | 21.7±4.3 | -29.2±0.6 | 22.4±2.4 |
| | *Spergularia* sp. | Vegetation | Spring | 5 | 11.2±3.9 | -27.5±1.9 | 20.9±2.4 |
| | | | Fall | 6 | 10.3±1.4 | -27.1±0.8 | 21.5±4.6 |
| | *Malva* spp. | Vegetation | Spring | 6 | 10.4±2.9 | -28.4±2.2 | 23.5±2.1 |
| | | | Fall | 6 | 8.2±0.9 | -29.1±1.3 | 25.1±4.5 |
| | *Plantago coronopus* | Vegetation | Spring | 5 | 16.0±1.5 | -29.1±1.3 | 23.9±3.3 |
| | | | Fall | 6 | 13.8±2.6 | -26.9±0.8 | 21.5±3.9 |
| | All Plants | | Spring | 22 | 12.0±3.2 | -28.4±1.7[a] | 23.7±3.0[a] |
| | | | Fall | 24 | 13.5±5.8 | -28.1±1.4[a] | 22.6±4.0[a] |
| Insect | *Coleoptera* larvae | Whole | Spring | 5 | 5.7±1.3 | -24.0±3.8 | 27.5±5.0 |
| | | | Fall | 6 | 5.9±1.0 | -24.6±1.2 | 24.7±1.6 |
| | *Farallonophilus cavernicolus* | Whole | Spring | 6 | 4.5±1.1 | -22.9±2.5 | 19.8±1.7 |
| | | | Fall | 6 | 4.8±0.6 | -18.9±0.8 | 23.8±2.0 |
| | *Oniscidea sp.* | Whole | Spring | 6 | 5.6±1.0 | -23.9±1.6 | 25.5±2.3 |
| | | | Fall | 6 | 5.2±0.6 | -21.0±0.6 | 23.7±3.0 |
| | *Araneae* spp. | Whole | Spring | 6 | 4.1±0.5 | -24.7±0.9 | 28.1±2.0 |
| | | | Fall | 6 | 4.0±0.4 | -23.6±1.2 | 26.8±1.7 |
| | All Insects | | Spring | 23 | 4.9±1.2 | -23.9±2.3[b] | 25.1±4.3[a] |
| | | | Fall | 24 | 5.0±1.0 | -22.0±2.4[c] | 24.7±2.4[a] |
| Intertidal | *Nucella emarginata* | Muscle | Spring | 6 | 3.7±0.2 | -11.7±2.0[d] | 14.8±1.4[b] |
| | | | Fall | 5 | 3.9±0.1 | -11.7±0.9[d] | 13.0±0.6[b] |
| Seabird | *Ptychoramphus aleuticus* | Egg membrane | Summer | 4 | 3.2±0.1 | -16.7±0.3 | 12.0±0.2 |
| | | Muscle | Summer | 4 | 3.0±0.1 | -16.2±0.5 | 16.1±1.7 |
| | *Larus occidentalis* | Egg membrane | Summer | 4 | 3.2±0.1 | -16.2±0.2 | 14.5±0.9 |
| | | Guano | Summer | 4 | 1.4±0.2 | -19.4±1.2 | 10.1±2.2 |
| | All Seabirds | All tissues | Summer | 16 | 2.7±0.8 | -17.1±1.5[e] | 13.2±2.7[b] |
