## Supplemental Table 3 for "Population Dynamics and Resource Availability Drive Seasonal Shifts in the Consumptive and Competitive Impacts of Introduced House Mice (*Mus musculus*) on an Island Ecosystem"

**Table S3**. The carbon ($\delta^{13}C$) and nitrogen ($\delta^{15}N$) stable isotope values, elemental concentration, and C/N ratio of major prey resource group averaged across all seasons on Southeast Farallon Island, CA in 2013 and used in stable isotope dietary mixing model analyses. Groups that share a superscript are not significantly different at the $P < 0.05$ level.

| Prey sources | *n* | Carbon % | Nitrogen % | C/N | $\delta^{13}C$ (‰) | $\delta^{15}N$ (‰) |
|---|---|---|---|---|---|---|
| Plant | 46 | 38.7±3.1 | 3.4±1.1 | 12.8±4.7 | $-28.2\pm1.5^{a}$ | $23.1\pm3.5^{a}$ |
| Insect | 47 | 43.7±8.5 | 14.9±2.7 | 4.9±1.0 | $-22.9\pm2.5^{b}$ | $24.9\pm3.5^{b}$ |
| Intertidal | 11 | 35.9±1.6 | 9.5±0.7 | 3.7±0.2 | $-11.7\pm1.5^{d}$ | $14.0\pm1.4^{c}$ |
| Seabird | 16 | 38.7±8.1 | 9.2±2.3 | 2.7±0.8 | $-17.1\pm1.5^{e}$ | $13.2\pm2.7^{c}$ |
| Salamander | 32 | 30.1±4.0 | 9.1±1.3 | 3.3±0.1 | $-20.7\pm1.4^{f}$ | $25.2\pm1.6^{b}$ |
